# Spinal cord extracts of amyotrophic lateral sclerosis spread TDP-43 pathology in cerebral organoids

**DOI:** 10.1101/2022.05.05.490760

**Authors:** Yoshitaka Tamaki, Jay P. Ross, Paria Alipour, Hélène Catoire, Daniel Rochefort, Makoto Urushitani, Ryosuke Takahashi, Joshua A. Sonnen, Stefano Stifani, Patrick A. Dion, Guy A. Rouleau

## Abstract

Amyotrophic lateral sclerosis (ALS) is a fatal neurodegenerative disorder caused by progressive loss of motor neurons and there is currently no effective therapy. Cytoplasmic mislocalization and aggregation of TAR DNA-binding protein 43 kDa (TDP-43) within the CNS is a pathological hallmark in sporadic ALS and prion-like propagation of pathogenic TDP-43 is thought to be implicated in disease progression. However, cell-to-cell transmission of pathogenic TDP-43 in the human CNS has not been confirmed experimentally.

Here we used induced pluripotent stem cells (iPSCs)-derived cerebral organoids as recipient CNS tissue model that are anatomically relevant human brain. We injected postmortem spinal cord protein extracts individually from three non-ALS or five sporadic ALS patients containing pathogenic TDP-43 into the cerebral organoids to validate the templated propagation and spreading of TDP-43 pathology in human CNS tissue.

We first demonstrated that the administration of spinal cord extracts from an ALS patient induced the formation of TDP-43 pathology that progressively spread in a time-dependent manner in cerebral organoids, suggesting that pathogenic TDP-43 from ALS functioned as seeds and propagated cell-to-cell to form de novo TDP-43 pathology. We also reported that the administration of ALS patient-derived protein extracts caused astrocyte proliferation to form astrogliosis in cerebral organoids, reproducing the pathological feature seen in ALS. Moreover, we showed pathogenic TDP-43 induced cellular apoptosis and that TDP-43 pathology correlated with genomic damage due to DNA double-strand breaks.

Thus, our results provide evidence that patient-derived pathogenic TDP-43 can mimic the prion-like propagation of TDP-43 pathology in human CNS tissue. Our findings indicate that our assays with human cerebral organoids that replicate ALS pathophysiology have a promising strategy for creating readouts that could be used in future drug discovery efforts against ALS.

## Introduction

Amyotrophic lateral sclerosis (ALS) is an adult-onset neurodegenerative disease characterized by progressive muscle weakness due to the selective loss of motor neurons in the nervous system, ultimately leading to death from respiratory failure. Approximately 10% of ALS cases have a positive family history (familial ALS), and the remaining 90% of cases occur sporadically (sporadic ALS).^1^ TAR DNA-binding protein 43 kDa (TDP-43) was discovered as a pathological hallmark of sporadic ALS and a subgroup of frontotemporal lobar degeneration (FTLD). TDP-43 functions as a DNA/RNA binding protein and is normally located in the nucleus. However, the misfolded form of TDP-43 accumulates in the cytosol as hyperphosphorylated TDP-43 aggregates in neurons and glia of the CNS in ALS and FTLD.^2,3^ In addition to TDP-43 pathology seen in ALS, the discovery of mutations in the gene encoding the TDP-43 protein in both sporadic and familial ALS suggests that TDP-43 protein is important in the pathogenesis of ALS.^4,5^

One of the key features of neurodegenerative diseases is the prion-like spreading of disease-related misfolded proteins such as tau in Alzheimer’s disease and α-synuclein in Parkinson’s disease.^6^ Accumulating evidence suggests that progression of neurodegenerative diseases is driven by the template-induced misfolding of endogenous normal proteins to pathological conformations with cell-to-cell transmission of disease-related proteins.^7^ The presence and propagation of such pathologically aggregated proteins leads to neuronal dysfunction and progression.^8-10^ The observation that ALS symptoms usually start in one region of the CNS and spreads to adjacent regions supports the notion that TDP-43 has prion-like properties. However, cell-to-cell propagation of pathogenic TDP-43 has been shown only in human cultured cells,^11-14^ not human tissues. Some groups have recently reported TDP-43 transmission in transgenic mice expressing human TDP-43, but not in wild type mice, administrated with tissue extracts from ALS patients.^15,16^ However, there is currently no study reporting the propagation of TDP-43 pathology in human CNS tissue.

Human induced pluripotent stem cells (iPSCs) are used to generate different disease models that can allow the study of disease pathogenesis.^17^ Moreover, iPSCs can be made to self-organize into three-dimensional cerebral organoids, which represents an anatomically and physiologically relevant model of human CNS tissue.^18^ Indeed, several studies using cerebral organoids studied disorders such as Parkinson’s disease,^19^ Alzheimer’s disease,^20^ and Creutzfeldt-Jakob disease.^21^ Yet, there are currently few studies of ALS using cerebral organoids.

Here we report that pathogenic TDP-43 extracted from postmortem ALS spinal cord tissue induces the formation of phosphorylated TDP-43 cytoplasmic aggregates in a time-dependent manner in human iPSC-derived cerebral organoids. We show that administration of pathogenic TDP-43 into cerebral organoids causes not only TDP-43 pathology but also astrogliosis, morphologically similar to what is seen in ALS postmortem tissue. We also show that pathogenic TDP-43 from ALS tissue facilitates cellular apoptosis and genomic damage. Our findings indicate that patient-derived pathogenic TDP-43 protein propagates in a prion-like manner and induces TDP-43 pathology in a time-dependent manner in a model that mimics human CNS environment.

## Materials and methods

### iPSCs cultures

Human iPSCs were generated from peripheral blood mononuclear cells (PBMCs) of consenting individuals. The PBMCs were reprogrammed into human iPSCs using the CytoTuneTM iPS Sendai reprogramming kit (Thermo Fisher Scientific) according to the manufacture’s protocol. iPSCs were cultured on a Matrigel (Corning) coated plate with mTeSR1 medium (STEMCELL Technologies) in 100% humidity incubator at 37°C under 5% CO_2_. iPSCs colonies were passaged at approximately 70-80% confluency. Human iPSCs information is listed in Supplementary Table 1.

### Cerebral organoid cultures

Cerebral organoids were differentiated from human iPSCs and cultured with STEMdiff Cerebral Organoid Kit (STEMCELL Technologies) according to the manufacture’s protocol. Once iPSCs colonies reached 70-80% confluency, iPSCs were placed in a 96-well round-bottom ultra-low attachment microplate (Corning) with EB formation medium (9,000 cells/well) containing ROCK inhibitor to generate embryoid bodies (EBs) from iPSCs, and EB formation medium was added every other day. At 5 days in vitro (DIV), EBs were moved to a 24-well ultra-low attachment plate (Corning) and cultured with induction medium for 48 hours. At 7 DIV, EBs were embedded into 15 μL of Matrigel and placed into a 6-well ultra-low attachment plate (Corning) with expansion medium for 72 hours to develop expanded neuroepithelia. At 10 DIV, expansion medium was replaced to maturation medium, and the 6-well cell culture plate was placed onto an orbital shaker with 65 rpm in a cell culture incubator to differentiate cells into cerebral organoids. Maturation medium was replaced every 3-4 days until they are used for experimental process.

### Human tissue

Postmortem sporadic ALS and non-ALS spinal cord frozen specimens were obtained from the Montreal Neurological Institute-Hospital (Montréal, Canada), the Douglas-Bell Canada Brain Bank (Montréal, Canada) and the London Neurodegenerative Diseases Brain Bank (London, UK). Tissue information is listed in Supplementary Table 2. Postmortem sporadic ALS-FTLD spinal cord formalin-fixed tissue for pathological diagnosis was collected from the Douglas-Bell Canada Brain Bank. Studies on human tissues were performed in accordance with the ethics board standards at McGill University (Montréal, Canada).

### Preparation of sarkosyl insoluble fractions from control and ALS spinal cords

The sarkosyl insoluble protein fraction from human postmortem frozen spinal cord was obtained by sequential protein extraction with increasing detergent strength buffers.^16^ Anterior horn from frozen cervical spinal cord was briefly sonicated and incubated with 5.0 mL/g high-salt buffer (10 mM Tris–HCl pH 7.4, 0.5 M NaCl, 2 mM EDTA, 10% sucrose, and 1 mM dithiothreitol) containing 1% Triton X-100 and protease and phosphatase inhibitor cocktails (Roche) (HS-TX buffer) for 30 minutes at 37°C. The protein extract was centrifuged at 25,000g for 30 minutes at 4°C and the supernatant was taken as HS-TX soluble fraction. The pellet was added and floated with cold HS-TX buffer containing 20% sucrose for 30 minutes on ice, and centrifuged at 25,000g for 30 minutes at 4°C for myelin removal. The remaining pellet was incubated with benzonaze (500 U/g) in high-salt buffer for 30 minutes on ice to remove tissue DNA and RNA. After centrifuging at 10,000g for 1 minute at 4°C, the pellet was incubated with 5.0 mL/g of high-salt buffer containing 2% sarkosyl (HS-2% sarkosyl) for 30 minutes at 37°C. The protein extract was centrifuged at 25,000g for 30 minutes at 22°C and the supernatant was taken as sarkosyl soluble fraction. The remaining pellet was washed twice in 3 mL/g cold PBS and centrifuged at 25,000g for 30 minutes at 15°C. The pellet was resuspended in 0.3 mL/g cold PBS and briefly sonicated, being taken as sarkosyl insoluble fraction. The protein concentrations of each fraction were measured with Bradford assay (Biorad).

### TDP-43 immunohistochemistry in postmortem human tissue

Standardized automated immunohistochemistry is performed using the Benchmark Ultra IHC platform with OptiView DAB IHC detection (Ventana Medical Systems) per manufacturer specification. Primary antibody (anti-phospho-TDP-43 (pS409); Cat#: TIP-PTD-P03, Cosmo Bio)^22^ is incubated at 1/1000 for 32 minutes at 36°C following antigen 40 minutes retrieval without amplification.

### Whole genome sequencing

DNA was extracted from a blood sample using standard salting out procedures. Whole genome sequencing (WGS) was performed at the Génome Québec Centre D’Expertise et de Services on a Novaseq 6000 sequencer (Illumina, Inc), using 150bp paired-end reads. All bioinformatics analyses were performed on the Béluga computing cluster of Compute Canada and Calcul Québec. An average genomic coverage of 39.31X was generated on the hg38 human reference genome. Alignment and variant calling were performed using the Illumina DRAGEN Bio-IT v3.8 platform.^23^ Variant filtration was performed using the Genome Analysis ToolKit (GATK) v4.1.8.1. Variants were annotated for functional consequence using VEP (ensembl version 98)^24^ and formatted using GEMINI.^25^ Only rare (gnomAD v3.0^26^ minor allele frequency less than 0.001) protein-altering variants within the coding regions of genes with previously reported association with ALS were considered.

### Injection of spinal cord tissue extracts into cerebral organoids

Cerebral organoids at 60 DIV were taken from maturation media and washed briefly with PBS at room temperature. 0.3 μL of sarkosyl-insoluble fraction of spinal cord extract was slowly injected into the cerebral organoids using a 32-gauge Hamilton Neurosyringe (Hamilton) at 1 mm depth. The injected cerebral organoids were placed back into maturation media and maintained in 100% humidity incubator at 37°C under 5% CO_2_.

### Protein extraction from cerebral organoids

After collected from maturation media and washed with cold PBS, cerebral organoids were briefly sonicated and lysed into 9x v/w cold RIPA buffer (50 mM Tris-HCl pH 7.4, 1 mM EDTA, 150 mM NaCl, 1% Nonidet-P40, 0.5% sodium deoxycholate, 0.1% sodium dodecyl sulfate) containing 1 mM phenylmethylsulfonyl fluoride (PMSF) and proteinase and phosphatase inhibitor cocktails (Roche). The protein extract was centrifuged at 25,000g for 30 minutes at 4°C, and the supernatant was used as RIPA-soluble fraction. The protein concentration of RIPA-soluble fraction was measured with Bradford assay (Biorad). The remaining pellet was washed with 5x v/w cold RIPA buffer and centrifuged at 25,000g for 30 minutes at 4°C. Subsequently, the pellet was resuspended and briefly sonicated in 4x v/w cold PBS, being taken as RIPA-insoluble fraction.

### Western blot analysis

Protein extract samples were eluted into 2x Laemmli sample buffer (Biorad) containing 2-mercaptethanol and denatured for 5 minutes at 95°C. The samples were loaded on a 12% TGX Stain-Free Fast Cast acrylamide gel (Biorad) and run for 30 minutes at 200 V. After electrophoresis, the protein samples were transferred onto a PVDF membrane (Millipore). The transferred PVDF membrane was blocked with 3% bovine serum albumin (BSA) in 0.1 M PBS/0.1% Tween 20 (0.1% PBS-T) at room temperature for 1 hour. The membrane was incubated with primary antibody diluted in 3% BSA/0.1% PBS-T at 4°C overnight. After washing with 0.1% PBS-T, the membrane was incubated with peroxidase-conjugated secondary antibody for 1 hour at room temperature. Targeted protein was detected by exposing the membrane to a mixture of the Clarity western ECL substrate (Biorad) and imaged using ChemiDoc MP Imager (Biorad).

### Cryogenic cerebral organoid section preparation

Cerebral organoids were immersed and fixed in 4% PFA at 4°C overnight and incubated in 30% sucrose/PBS at 4°C overnight. Subsequently the samples were embedded in 7.5% porcine skin gelatin solution with 10% sucrose/PBS after being incubated at 37°C for 1 hour, and frozen in liquid nitrogen. Frozen sections were cut into 14 μm thickness and mounted on glass slides. The frozen sections were stored at -80°C until experimental process.

### Immunofluorescence and microscopic analysis

After washing slides with PBS at 37°C for 10 minutes to remove gelatin from slides, cerebral organoid tissue was blocked with 5% BSA/0.1% PBS-T at room temperature for 1 hour. Tissue was reacted with primary antibody diluted in 5% BSA/0.1% PBS-T containing 0.05% sodium azide at 4°C overnight in a humidified chamber. After washing with 0.1% PBS-T, tissue was incubated with Alexa Fluor-conjugated secondary antibody at room temperature for 1 hour in the dark. After washing with 0.1% PBS-T, a coverslip was mounted with antifade mounting medium containing DAPI (Vector Laboratories). Images were obtained with Leica TCS SP8 confocal microscope (Leica) using the Leica LAS-X software.

### TUNEL stain assay

TUNEL stain assay was evaluated with BrdU-Red TUNEL assay kit (Abcam) according to the manufacture’s protocol. After removing gelatin from a frozen cerebral organoid section slide with PBS at 37°C for 10 minutes, cerebral organoid tissue was added with 20 μg/mL proteinase K solution (100 mM Tris-HCl pH 8.0 and 50 mM EDTA) at room temperature for 5 minutes, and subsequently mounted with 4% PFA at room temperature for 5 minutes. After washing the slide, the tissue was reacted with DNA labeling solution containing TdT enzyme and Br-dUTP at 37°C for 1 hour in a dark and humidified incubator. The tissue was next incubated with anti-BrdU-Red antibody at room temperature for 30 minutes in the dark. After washing the slide, a coverslip was mounted with antifade mounting medium containing DAPI (Vector Laboratories). Images were obtained with Leica TCS SP8 confocal microscope using the Leica LAS-X software.

### Statistical analysis

Statistical analysis definitions were described in each figure legends and calculated using GraphPad Prism 7 software. Unpaired *t*-test was performed when two groups were compared, and two-way ANOVA with Tukey’s multiple comparisons test was used when multiple groups were compared in different conditions. A *P* value of < 0.05 was considered statistically significant.

### Antibodies

Antibodies used in this study are listed in the Supplementary Table 3.

### Ethics

Human PBMCs were collected to generate iPSCs after obtaining written informed consent forms in accordance with the institutional human ethics committee guidelines. All the experimental procedures using postmortem human tissue samples were approved by McGill University Health Centre Research Ethics Board (IRB00010120).

### Data availability

The authors confirm that the data supporting the findings of this study are available within the article upon reasonable request.

## Results

### Human iPSCs differentiate into cerebral organoids

Based on work by Lancaster *et al*.^27^ we can generate an *in vitro* model of the developing human brain with iPSCs. We first differentiated iPSCs into cerebral organoids to create human CNS model that we could inoculate with pathogenic TDP-43. At 60 DIV, our cerebral organoids grew to over 1 mm in diameter with CNS-like tissue organization that expressed TUJ1, a neural marker (Supplementary Fig. 1A). Immunofluorescence analysis showed that cerebral organoids mainly expressed PAX6-positive neural progenitor cells at 30 DIV. By 60 DIV they had many CTIP2-positive cells, mostly in the surface (Supplementary Fig. 1B), as well as TUJ1-positive cells, indicating that they contained neural cortical cells (Supplementary Fig. 1C). In addition, these cerebral organoids simultaneously expressed CTIP2-positive cells and SOX2-positive neural progenitor cells (Supplementary Fig. 1C). These data demonstrated that the human iPSCs were successfully differentiated into cerebral organoids, recapitulating to some extent human brain tissue.

### ALS patient-derived protein extracts induce cytoplasmic pTDP-43 aggregates

To investigate the intracellular seeding phenomenon of pathogenic TDP-43 in recipient cerebral organoids, we used sporadic ALS spinal cord protein extracts which contained pathogenic phosphorylated TDP-43 (pTDP-43) at Ser409/Ser410 sites (pS409/410) observed in ALS.^22^ We collected postmortem spinal cord tissues from five sporadic ALS patients including one with the chromosome 9 open reading frame 72 (*C9orf72*) repeat expansion mutation and from three non-ALS cases as negative controls (Supplementary Table 2). The spinal cord protein extracts were separated into detergent-soluble or -insoluble fractions with increasing detergent strength buffers using previously described procedures.^16^ Western blot analysis showed that all sarkosyl-insoluble protein extracts from five sporadic ALS patients contained both full-length and C-terminal fragments (CTFs) of pTDP-43 (pS409/410), consistent with the biochemical profile of pathological TDP-43 in sporadic ALS (Supplementary Fig. 2).^2,3,22^

To determine if patient-derived pathogenic TDP-43 propagates in our human CNS tissue model, sarkosyl-insoluble protein extracts containing pTDP-43 were administrated as pathogenic seeds into mature cerebral organoids at 60 DIV through a fine needle injection. We used cerebral organoids differentiated from two human iPSCs lines in this study; AJC001 iPSCs line derived from non-ALS case and TD17 from the patient who was clinically and pathologically diagnosed with sporadic ALS-FTLD without mutations in any ALS-related genes (Supplementary Table 1 and Supplementary Fig. 3). Although the whole genome sequencing showed that TD17 iPSCs line had a GLE1 p.C39Y variant that has not previously been reported in ALS and it was predicted *in silico* to be benign.

At 8 weeks post injection (p.i.) with ALS patient-derived protein extract, AJC001 cerebral organoids had few pTDP-43 aggregates (Fig. 1A). In contrast, TD17 cerebral organoids at 8 weeks p.i. had a larger number of pTDP-43 aggregates in the neural cytoplasm (Fig. 1B). Considering that injection of non-ALS case-derived protein extracts did not result in the formation of pTDP-43 aggregates in TD17 cerebral organoids, pTDP-43 aggregates were not derived from the cerebral organoids themselves but were induced by the administration of protein extracts containing pTDP-43. In addition, pTDP-43 aggregates were observed not only in TUJ1-positive neural cells but also in glial fibrillary acidic protein (GFAP)-positive astrocytes (Supplementary Fig. 4). The formation of pTDP-43 aggregates and their distribution in cerebral organoids resembles TDP-43 pathology seen in ALS.

**Figure 1.**
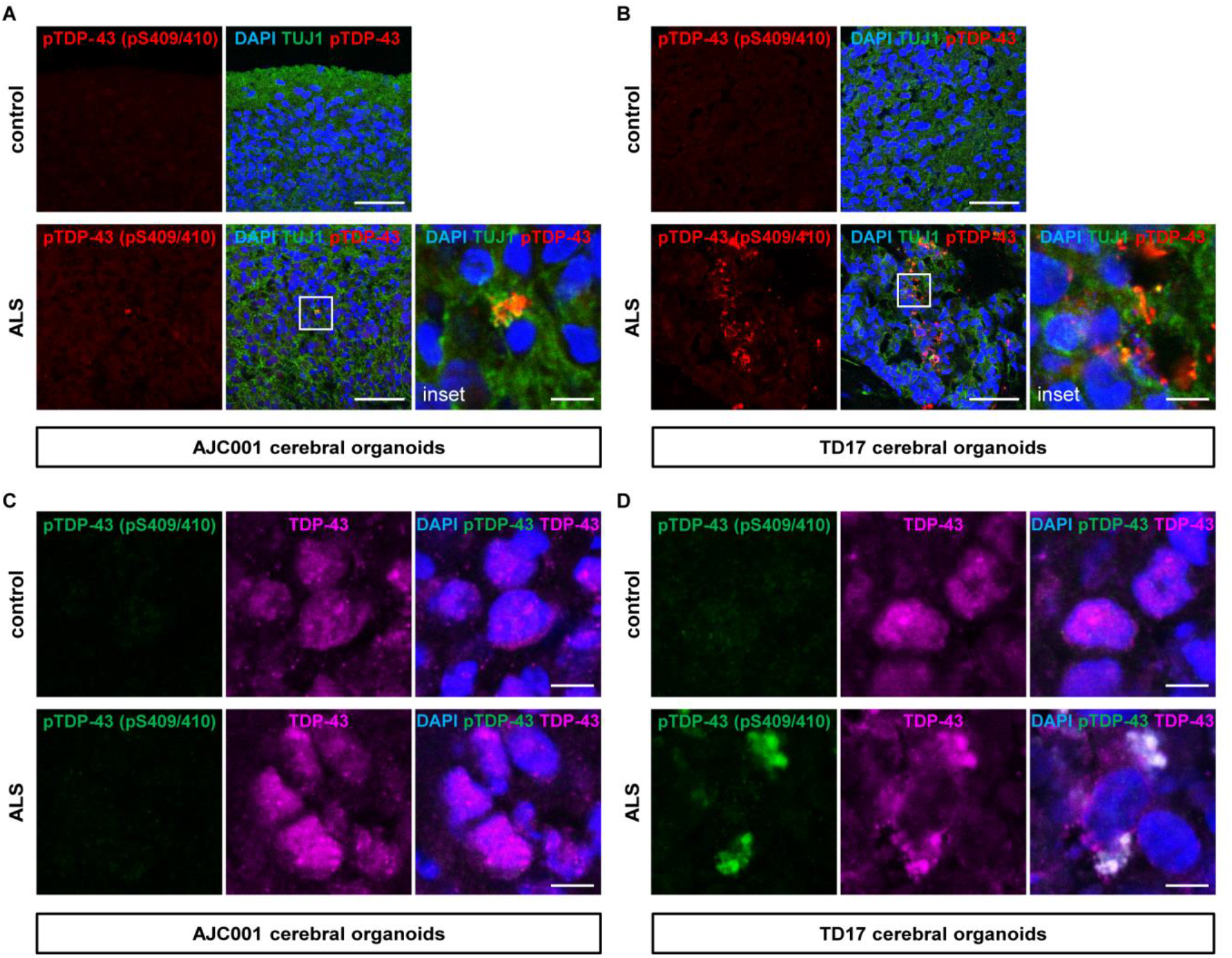
ALS patient-derived protein extracts induce the formation of TDP-43 pathology. (**A** and **B**) Double-labelled immunofluorescence images of pTDP-43 (pS409/410) and TUJ1 staining AJC001 (**A**) or TD17 cerebral organoids (**B**) at 8 weeks p.i. of protein extract from control (control 1) or ALS (patient 4). A higher magnification of the white line box is shown in the right inset. Scale bars = 50 μm (10 μm for insets). (**C** and **D**) Immunofluorescence image of AJC001 (**C**) or TD17 cerebral organoids (**D**) double-labelled with pTDP-43 (pS409/410) and TDP-43 at 8 weeks p.i. of spinal cord extract from control (control 1) or ALS (patient 3). Scale bars = 10 μm. DAPI was used for counterstaining of nuclei.

TDP-43 is an RNA/DNA binding protein that localizes predominantly in the nuclei. However, in most sporadic ALS patients pathological pTDP-43 aggregates can be found in the cytosol of neurons and glia cells with loss of nuclear TDP-43.^1-3^ We next examined TDP-43 distribution in cerebral organoids at 8 weeks p.i.. TDP-43 was diffusely distributed in the nuclei of the AJC001 cerebral organoids injected with either control or ALS patient-derived protein extract (Fig. 1C). However, in TD17 cerebral organoids injected with ALS patient-derived protein extract, cells had cytoplasmic aggregates and showed reduced nuclear TDP-43, recapitulating TDP-43 pathology (Fig. 1D). These data suggest that the administration of pathogenic TDP-43 into ALS-derived cerebral organoids induced loss of nuclear TDP-43 function as well as pathogenic pTDP-43 localization to the cytosol.

### ALS patient-derived protein extracts induce spread of TDP-43 pathology in cerebral organoids

In TDP-43-associated FTLD and sporadic ALS, propagation of pathogenic TDP-43 has been demonstrated in adherent human cultured cells,^11-14^ and recently in transgenic mice.^15,16^ Based on these reports, we wished to determine if patient-derived pathogenic TDP-43 could seed and induce the templated self-propagation of TDP-43 pathology in human cerebral organoids.

Immunofluorescence analysis of AJC001 cerebral organoids at 2, 4, and 8 weeks p.i. with ALS patient-derived protein extract had no formation of pTDP-43 aggregates at either 2 or 4 weeks p.i., and minor amounts of pTDP-43 aggregates at 8 weeks p.i. (Fig. 2A). Analysis of same treatment of TD17 cerebral organoids showed pTDP-43 aggregates at 2 weeks p.i., and the amount of aggregates increased at 4 and 8 weeks p.i. (Fig. 2B). To further investigate the development of TDP-43 pathology over time, we measured detergent-insoluble pTDP-43 in cerebral organoids at 2, 4, and 8 weeks p.i. using Western blot analysis. In AJC001 cerebral organoids, RIPA-insoluble pTDP-43 were hardly detectable at 2 and 4 weeks p.i., and slightly detectable at 8 weeks p.i. (Fig. 2C). In contrast, RIPA-insoluble full-length pTDP-43 as well as CTFs in TD17 cerebral organoids were detectable at 2 weeks p.i. and they increased in quantity over time to 8 weeks p.i. (Fig. 2D), which was consistent with the immunofluorescence results. Interestingly, these results were seen with all 5 different ALS patient-derived protein extracts, albeit with different intensities.

**Figure 2.**
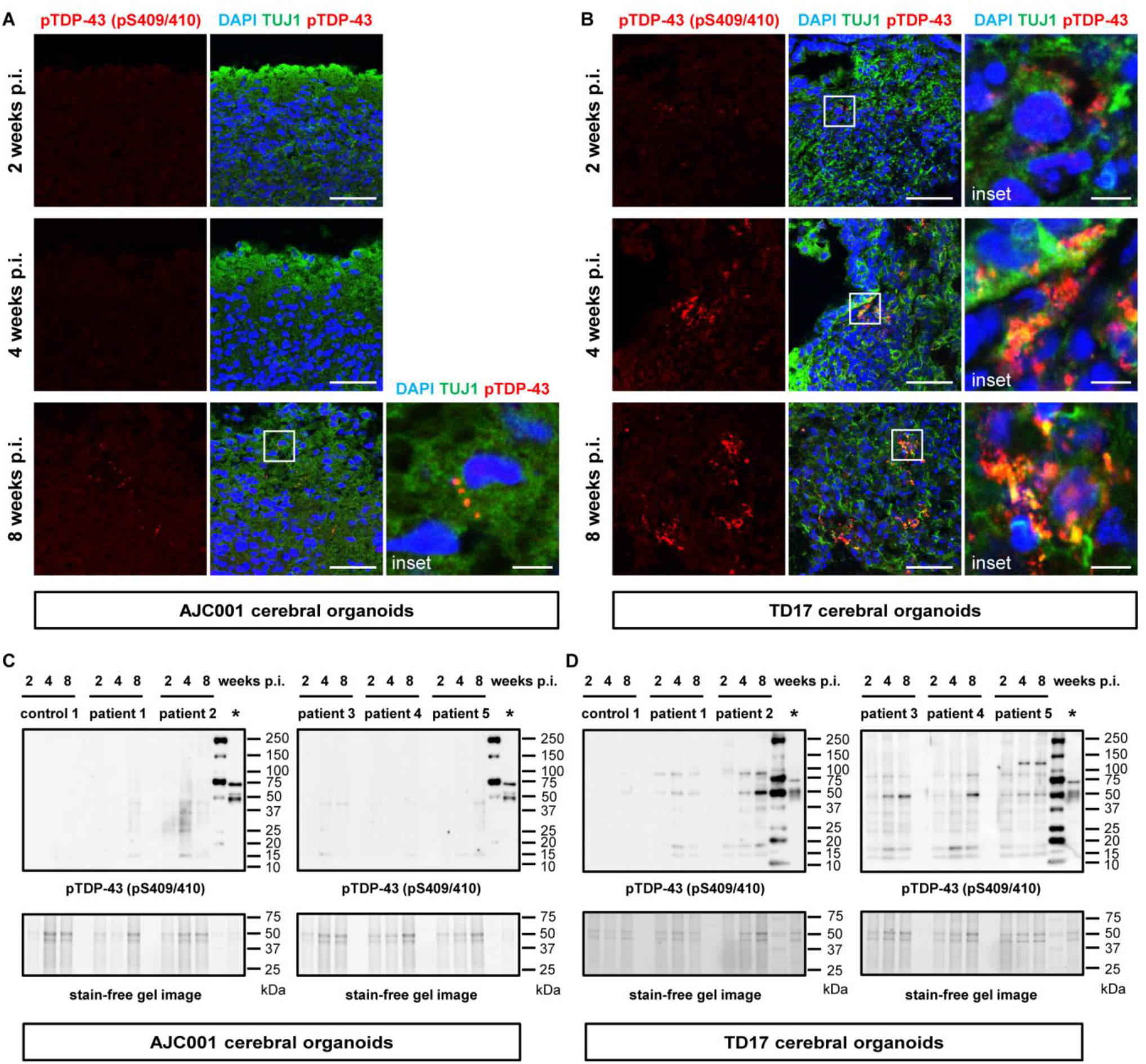
TDP-43 pathology spreads in a time-dependent manner in cerebral organoids. (**A** and **B**) Immunofluorescence images of pTDP-43 (pS409/410) and TUJ1 staining AJC001 (**A**) or TD17 cerebral organoids (**B**) at 2, 4 and 8 weeks p.i. of protein extract from ALS (patient 5). A higher magnification of each white line box is shown in the right inset. DAPI was used for counterstaining of nuclei. Scale bars = 50 μm (10 μm for insets). (**C** and **D**) Western blot analysis of RIPA-insoluble protein fractions of AJC001 (**C**) or TD17 cerebral organoids (**D**) at 2, 4 and 8 weeks p.i. of protein extract from control (control 1) or individual ALS cases (patient 1-5), immunoblotted with pTDP-43 (pS409/410) antibody. Sarkosyl-insoluble fraction of spinal cord from patient 5 was loaded on the lanes marked with asterixis (*) as a positive control for pTDP-43 immunoblots. The second right lanes were loaded with a protein marker. Stain-free gel images were used for protein loading controls.

A G_4_C_2_ repeat hexanucleotide expansion in the *C9orf72* gene is reported as the most common genetic cause of ALS and FTLD.^28,29^ A glycine-arginine repeat dipeptide (GR repeat) translated from expanded G_4_C_2_ repeat is thought to play a crucial role in inducing TDP-43 pathology through the disruption of nucleocytoplasmic transport and stress granule dynamics.^30-32^ Previous reports show that dipeptide repeats including GR repeats translated from the expanded *C9orf72* gene have a potential to transmit cell-to-cell via exosome-dependent and exosome-independent pathways.^33^ Moreover, the GR repeat promotes aggregation of endogenous TDP-43, mediating sequestration of full-length TDP-43 to induce cytoplasmic TDP-43 inclusion formation.^34^ Consistent with these reports, GR repeat proteins were seen in TD17 cerebral organoids injected with protein extracts from *C9orf72* gene-expanded ALS (C9orf72-ALS), but not from non-C9orf72-ALS. In addition, pTDP-43 aggregates colocalized with GR repeat proteins in TD17 cerebral organoids injected with protein extracts from C9orf72-ALS, suggesting that GR repeat protein spread and recruited the formation of TDP-43 pathology in cerebral organoids that do not have the *C9orf72* gene expansion (Supplementary Fig. 5). This result reproduces the C9orf72 pathology where GR repeat protein colocalizes with pathological TDP-43 inclusions in the motor cortex of C9orf72-ALS patient.^35^

Taken together, our results indicated that ALS patient-derived protein extract containing pTDP-43 were pathogenic seeds that spread TDP-43 pathology in cerebral organoids, especially cerebral organoids from a patient who had ALS, recapitulating the pathogenic TDP-43 propagation phenomenon in human CNS tissue.

### ALS patient-derived protein extracts cause astrogliosis in cerebral organoids

Astrocytes are key components of the CNS that are involved in multiple neural homeostatic functions. They influence synaptic function and formation, regulate the concentration of neurotransmitters at the synapse, supply metabolites to neurons, provide neurotrophic factors, and aid in the repair of damaged neural tissue.^36-38^ ALS pathology displays astrogliosis correlated to astrocyte proliferation and hypertrophy in the CNS, including spinal cord, cerebral cortex, and subcortical white matter.^39^ Therefore, we investigated whether the formation of TDP-43 pathology is accompanied by astrogliosis in cerebral organoids. Immunofluorescence analysis showed that, while AJC001 cerebral organoids did not show GFAP-positive astrocyte proliferation, TD17 cerebral organoids exhibited an increasing amount of astrocytes after they were injected with ALS patient-derived protein extracts (Fig. 3A and B). The statistical analysis of the immunofluorescence signal demonstrated that GFAP-positive astrocytes were significantly proliferated in TD17 cerebral organoids that were injected with ALS patient-derived protein extracts (Fig. 3C). To confirm the immunofluorescence results, we further investigated the expression of GFAP in cerebral organoids using Western blot analysis. While the amount of GFAP expression was not different between AJC001 cerebral organoids injected with control or ALS patient-derived protein extract, a significant difference in GFAP expression was seen in TD17 cerebral organoids (Fig. 3D and E). Densitometry analysis of Western blot results confirmed this finding (Fig. 3F and G). These data demonstrate that ALS patient-derived protein extracts containing pTDP-43 induced astrogliosis as well as TDP-43 pathology in cerebral organoids of an ALS patient.

**Figure 3.**
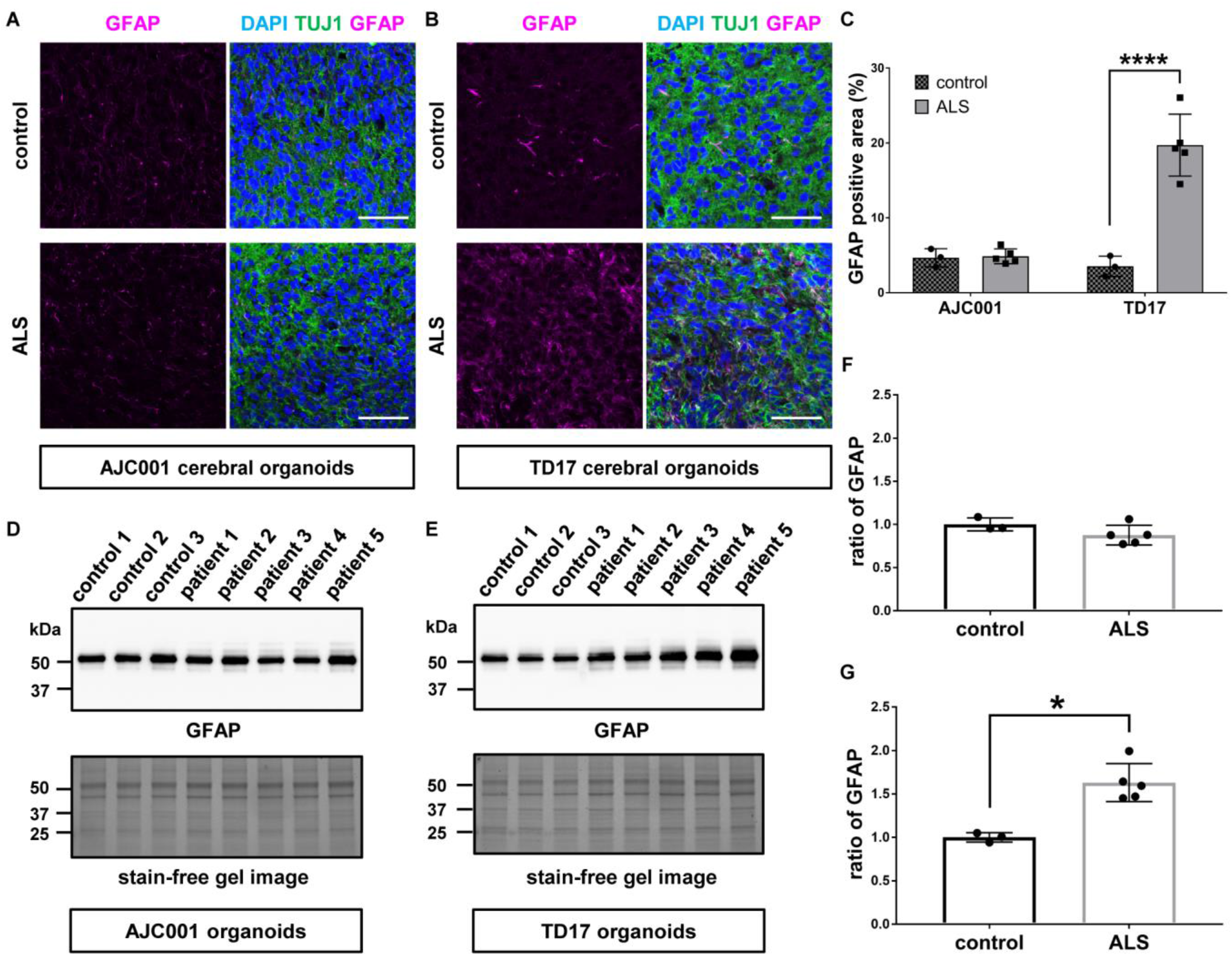
ALS patient-derived protein extract injection induces astrogliosis in cerebral organoids. (**A** and **B**) Immunofluorescence images of GFAP and TUJ1 staining AJC001 (**A**) or TD17 cerebral organoids (**B**) at 8 weeks p.i. of protein extract from control (control 1) or ALS (patient 3). DAPI was used for counterstaining of nuclei. Scale bars = 50 μm. (**C**) Quantification analysis of immunofluorescence images of GFAP staining AJC001 and TD17 cerebral organoids at 8 weeks p.i. of protein extract from individual controls (control 1-3) or ALS cases (patient 1-5) represented in (**A**) and (**B**). Percentages of GFAP-positive area were measured with ImageJ software. Bar plots show mean ± standard deviation (SD) with individual points representing a different proteins extract. Differences were evaluated by two-way ANOVA with Tukey’s multiple comparisons test. \*\*\*\**p*<0.001. (**D** and **E**) Western blot analysis of GFAP expression in AJC001 (**D**) or TD17 cerebral organoids (**E**) at 8 weeks p.i. of protein extracts from individual controls (control 1-3) or ALS cases (patient 1-5). A stain-free gel image was used for protein loading controls. (**F** and **G**) Densitometric quantification of Western blot analysis in (**D**) and (**E**), respectively, using ImageJ software. Each data point was obtained by normalization to bands in stain-free gel blot images. Bar plots show mean ± SD with individual points representing a different protein extract from controls (control 1-3) and ALS (patient 1-5). Differences were evaluated by unpaired *t*-test. \**p*<0.05.

### ALS patient-derived protein extracts induce cellular apoptosis and DNA double-strand breaks

Recent studies show that TDP-43 pathology correlates with neural apoptosis due to DNA double-strand break repair defects.^40-42^ Thus, we next examined the cytotoxic effects of TDP-43 pathology and DNA damage in cerebral organoids injected with ALS patient-derived protein extracts.

We first performed TUNEL stain assay to evaluate cellular apoptosis in cerebral organoids. Few TUNEL-positive cells were seen in AJC001 cerebral organoids after the administration of protein extracts from either control or ALS patient (Fig. 4A). However, TD17 cerebral organoids had several TUNEL-positive cells after injection with ALS patient-derived protein extracts (Fig. 4B). Statistical analysis confirmed the increase in TUNEL-positive cells in TD17 cerebral organoids injected with ALS patient-derived protein extract (Fig. 4C). We next investigated the expression of cleaved caspase-3 protein, a marker of cellular apoptosis. In immunofluorescence analysis, AJC001 cerebral organoids had little cleaved caspase-3, whether they were injected with control or ALS patient-derived protein extracts (Fig. 4D). On the other hand, the expression of cleaved caspase-3 was increased in TD17 cerebral organoids after injection with ALS patient-derived protein extracts (Fig. 4E). Statistical analysis confirmed that cleaved caspase-3 was significantly higher in TD17 cerebral organoids that received ALS patient-derived protein extracts (Fig. 4F). Using Western blot analysis, we further investigated the amount of cleaved caspase-3 expressions in our models. As was found by immunofluorescence, there was no difference in cleaved caspase-3 expression levels in AJC001 cerebral organoids injected either with control or ALS patient-derived protein extracts, while in TD17 cerebral organoids there was more cleaved caspase-3 expression with ALS patient-derived protein extracts (Fig. 4G and H). The statistical evaluation using densitometry in Western blot analysis also revealed that cleaved caspase-3 expression was the same in AJC001 cerebral organoids injected with control or ALS patient-derived protein extract, but it was significantly increased in TD17 cerebral organoids injected with ALS patient-derived protein extracts (Fig. 4I and J). These results indicate that ALS patient-derived protein extracts containing pathogenic TDP-43 trigger neural apoptosis in ALS-derived cerebral organoids.

**Figure 4.**
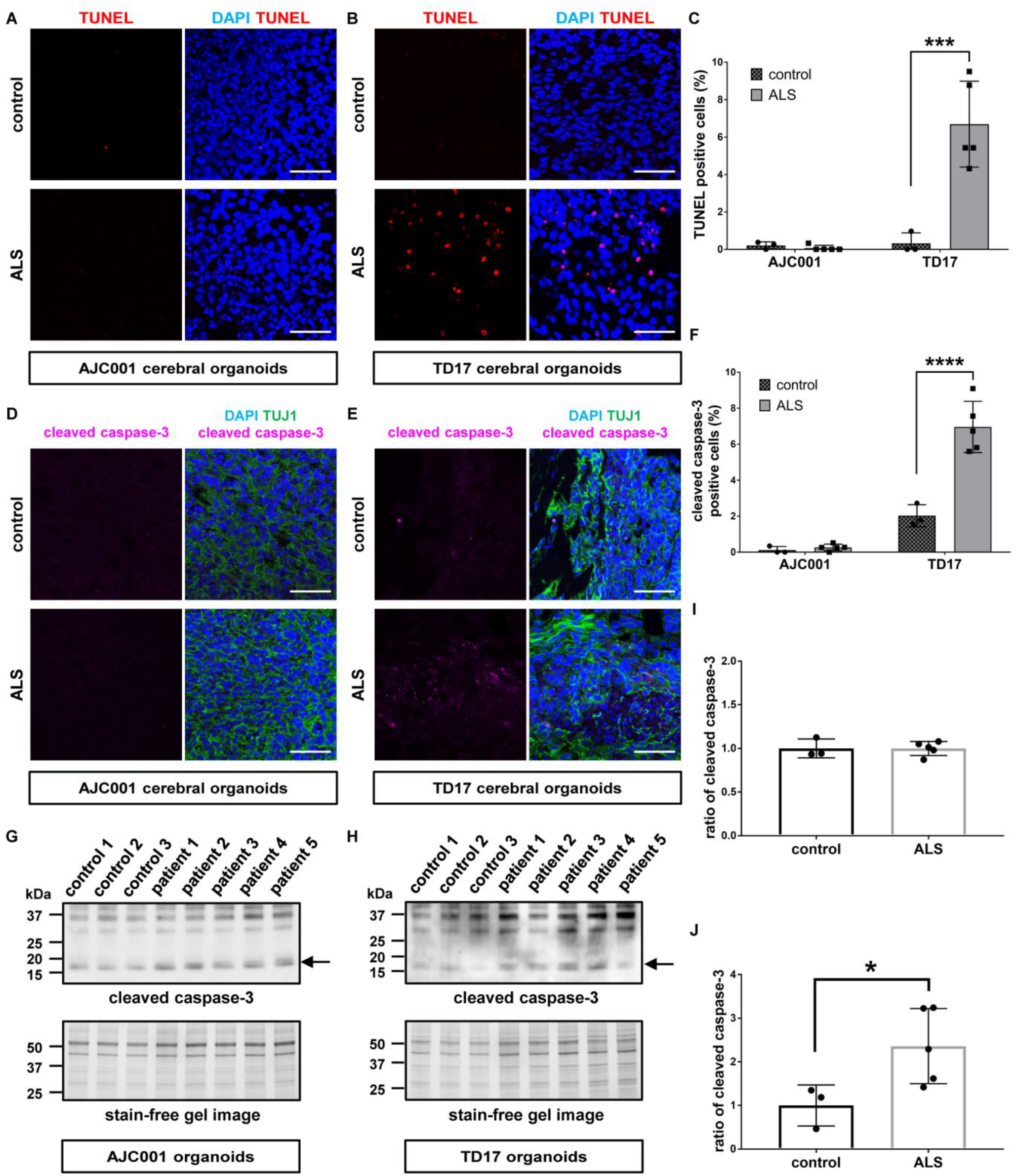
ALS patient-derived protein extracts cause cellular apoptosis in cerebral organoids. (**A** and **B**) TUNEL stain images of AJC001 (**A**) or TD17 cerebral organoids (**B**) at 8 weeks p.i. of protein extract from control (control 1) or ALS (patient 4). DAPI was used for counterstaining of nuclei. Scale bars = 50 μm. (**C**) Quantification analysis of TUNEL stain images of AJC001 and TD17 cerebral organoids at 8 weeks p.i. of protein extract from individual controls (control 1-3) or ALS cases (patient 1-5) representatively shown in (**A**) and (**B**). Bar plots show mean ± SD with individual points representing a different protein extract. Percentages of TUNEL-positive cells were normalized to number of DAPI-positive cell nuclei. Differences were evaluated by two-way ANOVA with Tukey’s multiple comparisons test. \*\*\**p*<0.005. (**D** and **E**) Immunofluorescence images of cleaved caspase-3 and TUJ1 staining AJC001 (**D**) or TD17 cerebral organoids (**E**) at 8 weeks p.i. of protein extract from control (control 1) or ALS (patient 5). Scale bars = 50 μm. (**F**) Quantification analysis of cleaved caspase-3 staining immunofluorescence images of AJC001 and TD17 cerebral organoids at 8 weeks p.i. of protein extract from individual controls (control 1-3) or ALS cases (patient 1-5) represented in (**D**) and (**E**). Bar plots show mean ± SD with individual points representing a different protein extract. Percentages of cleaved caspase-3-positive cells were normalized to number of DAPI-positive cell nuclei. Differences were evaluated by two-way ANOVA with Tukey’s multiple comparisons test. \*\*\*\**p*<0.001. (**G** and **H**) Western blot analysis of cleaved caspase-3 expression in AJC001 (**G**) or TD17 cerebral organoids (**H**) at 8 weeks p.i. of protein extracts from individual controls (control 1-3) or ALS (patient 1-5). A stain-free gel image was used for protein loading controls. Arrows indicate cleaved caspase-3 protein bands detected in approximately 17 kDa molecular weight. (**I** and **J**) Densitometric quantification of Western blot analysis in (**G**) and (**H**), respectively, using ImageJ software. Bar plots show mean ± SD with individual points representing a different protein extract. Each data point was obtained by normalization to bands in stain-free gel blot images. Differences were evaluated by unpaired *t*-test. \**p*<0.05.

Recent studies uncovered the relationship between TDP-43 pathology and DNA damage which causes neural apoptosis. They reported that pathological TDP-43 induces DNA double-strand breaks (DSBs) repair defects in ALS.^41,42^ DSBs evoke phosphorylation of histone variant protein H2AX on serine 139.^43^ The phosphorylated H2AX protein, gamma-H2AX (γH2AX), forms nuclear foci which recruit DNA repair proteins. Hence the formation of γH2AX nuclear foci is widely used as a specific marker to evaluate DSBs.^44^ Based on these previous reports, we investigated by immunofluorescence whether the cellular apoptosis caused by pathogenic TDP-43 in cerebral organoids was associated with DNA damage by detecting γH2AX. AJC001 cerebral organoids showed only a few γH2AX nuclear foci whether they were injected with extracts from control or ALS patient-derived protein extracts (Fig. 5A). In contrast, TD17 cerebral organoids had more γH2AX nuclear foci after they were injected with ALS patient-derived protein extract than from control protein extract (Fig. 5B). Moreover, some γH2AX colocalized with cytoplasmic pTDP-43 aggregates in TD17 cerebral organoids, suggesting that pathogenic pTDP-43 recruited DSBs repair proteins with γH2AX in the cytosol (Fig. 5B). Statistical analysis confirmed these findings (Fig. 5C). We next performed Western blot analysis to measure the expression of γH2AX. Expression levels were not different in AJC001 cerebral organoids between controls and ALS patient-derived protein extracts, and expression was slightly higher in TD17 cerebral organoids injected with ALS patient-derived protein extracts than in the control injected organoids (Fig. 5D and E). The densitometry of Western blot results showed that, while there was no difference for the ratio of γH2AX expression between controls and ALS in AJC001 cerebral organoids, injection of ALS patient-derived protein extracts in TD17 cerebral organoids significantly enhanced γH2AX expression compared to controls (Fig. 5F and G).

Overall, our data demonstrates that pathogenic TDP-43 from sporadic ALS patient-derived protein extracts has the potential to seed and propagate in ALS-derived cerebral organoids used as human CNS tissue model, inducing astrogliosis and cellular apoptosis concomitant with DSBs.

**Figure 5.**
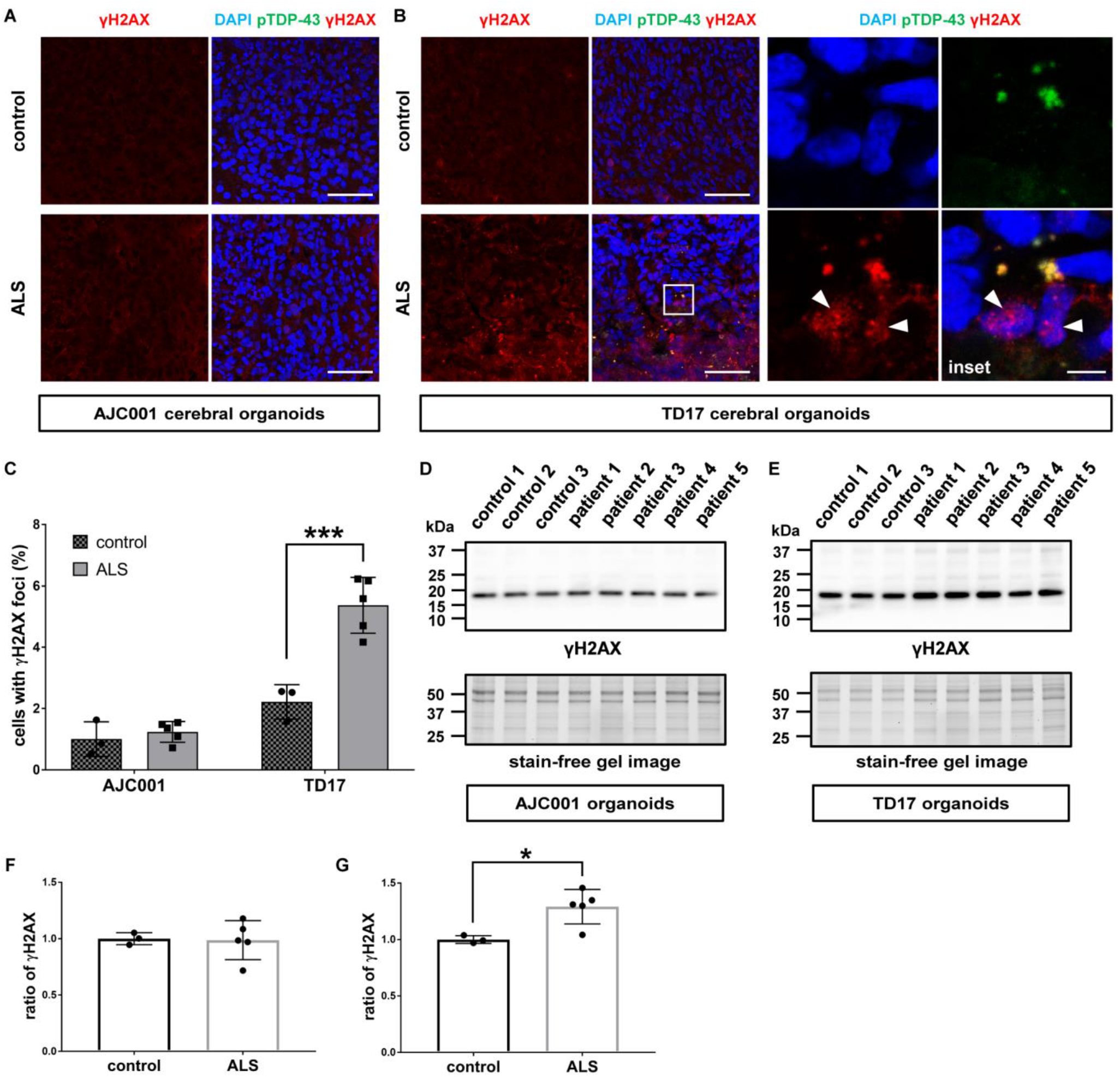
TDP-43 pathology promotes DNA damage in cerebral organoids. (**A** and **B**) Immunofluorescence images of γH2AX and pTDP-43 (pS409/410) staining AJC001 (**A**) or TD17 cerebral organoids (**B**) at 8 weeks p.i. of protein extract from control (control 1) or ALS (patient 2). A higher magnification of white line box is shown in the right inset. DAPI was used for counterstaining of nuclei. Arrow heads indicate γH2AX nuclear foci. Scale bars = 50 μm (10 μm for inset). (**C**) Quantification analysis of γH2AX staining immunofluorescence images of AJC001 and TD17 cerebral organoids at 8 weeks p.i. of protein extract from individual controls (control 1-3) or ALS cases (patient 1-5) represented in (**A**) and (**B**). Bar plots show mean ± SD with individual points representing a different protein extract. Percentages of cells harboring γH2AX nuclear foci were normalized to number of DAPI-positive cell nuclei. Differences were evaluated by two-way ANOVA with Tukey’s multiple comparisons test. \*\*\**p*<0.005. (**D** and **E**) Western blot analysis of γH2AX expression in AJC001 (**D**) or TD17 cerebral organoids (**E**) at 8 weeks p.i. of protein extracts from individual controls (control 1-3) or ALS (patient 1-5). A stain-free gel image was used for protein loading controls. (**F** and **G**) Densitometric quantification of Western blot analysis in (**D**) and (**E**), respectively, using ImageJ software. Bar plots show mean ± SD with individual points representing a different protein extract from controls (control 1-3) and ALS (patient 1-5). Each data point was obtained by normalization to bands in stain-free gel blot images. Differences were evaluated by unpaired *t*-test. \**p*<0.05.

## Discussion

We demonstrate that ALS patient-derived protein extracts containing pathogenic TDP-43 have the potential to spread cell-to-cell and to form phosphorylated TDP-43 cytoplasmic aggregates replicating ALS pathology in a time-dependent manner in recipient human cerebral organoids. We also showed that the TDP-43 pathology causes astrogliosis, cellular apoptosis and genomic damage. Our findings support the recent reports describing that pathogenic TDP-43 show prion-like propagation in murine brain and spinal cord of TDP-43 transgenic animal administrated with ALS patient-derived tissue^15,16^ and that TDP-43 pathology correlates with DSBs *in vitro* and *in vivo*.^41,42^ These results were relatively consistently seen with ALS patient-derived protein extracts from five different individuals. Ours is the first study to elucidate the seeding activity of patient-derived pathogenic TDP-43 and the formation of TDP-43 pathology in a human CNS tissue model.

We use mature iPSCs-derived cerebral organoids as a model of the human brain. These organoids have a complex three-dimensional structure with a wide variety of neuronal and glial cell types, so resembles human brain tissue.

Interestingly, the cerebral organoids derived from sporadic ALS-FTLD patient (TD17 cerebral organoids), but not cerebral organoids from a control individual, showed increased cytoplasmic pTDP-43 aggregates and the TDP-43 pathology as early as two weeks after the administration of ALS patient-derived protein extracts. Cerebral organoids differentiated from an unaffected healthy control (AJC001 cerebral organoids) formed small amounts of cytoplasmic pTDP-43 accumulation only eight weeks after the injection. Our findings raise the possibility that cells from someone who developed ALS carry some factors making them more susceptible to TDP-43 induced pathology. All known genetic factors have been ruled out as a possible cause for this susceptibility. Perhaps some unknown genetic factors, or some residual epigenetic marks, are responsible for this susceptibility to prion-like spread of pathogenic TDP-43. Based on the experiments for TDP-43 propagation in transgenic mice, Porta *et al*.^16^ describe that both concentration and subcellular localization of TDP-43 protein are important for TDP-43 aggregation and that disturbances in nuclear-cytoplasmic TDP-43 protein homeostasis may play a role in TDP-43 nucleation and aggregation. Although our cerebral organoids from iPSCs derived from an ALS-FLTD patient do not exhibit apparent cytoplasmic TDP-43 expression before the administration of ALS patient-derived protein extract, they may have a defect in nuclear-cytoplasmic TDP-43 translocations that facilitates TDP-43 aggregation once exposed to pathogenic TDP-43 from ALS.

Our results showing the time-dependent increase of pTDP-43 aggregates in recipient cerebral organoids administrated with ALS patient-derived protein extracts suggests that exogenous pathogenic TDP-43 acts as seeds and propagates cell-to-cell to form de novo TDP-43 pathology by altering normal endogenous TDP-43 to pathological conformations. Although the specific mechanisms for prion-like propagation of pathogenic TDP-43 remains enigmatic, accumulating evidence indicates that exosomes play a role in transferring several pathological proteins associated with neurodegenerative disorders such as tau, α-synuclein, and SOD1.^45-47^ As for TDP-43, a previous report demonstrates that secreted exosomes from ALS brain causes cytoplasmic TDP-43 distribution in neural cultured cells, suggesting that exosomes may contribute to propagation of TDP-43 pathology.^48^ Although it remains unclear whether exosome-mediated transfer is a major pathway for TDP-43 propagation in ALS, our study using cerebral organoids and patient-derived protein extract has the potential to be a useful model for evaluating molecular factors that cause TDP-43 transmission.

Astrogliosis characterized by hypertrophy and proliferation of astrocytes as well as upregulation of GFAP expression is one of the key features observed in ALS pathology.^39^ Our model successfully reproduces the astrogliosis seen in ALS. Of note, astrocytes are involved in both familial and sporadic ALS pathogenesis by releasing various toxic factors that provoke neuroinflammation and neurotoxicity.^49-51^ The increase in activated astrocytes in our ALS cerebral organoid model may contribute to cellular death and spread of misfolded TDP-43 via secreted neurotoxic molecules.

We show here that the administration of ALS patient-derived protein extracts also induces cellular apoptosis and DSBs in recipient cerebral organoids, suggesting a link between apoptotic cellular death and genomic damage in TDP-43 pathology. Genome damage-mediated neuronal apoptosis in ALS has been reported in several studies. TDP-43 is a critical component of the DSBs repair pathway and physiologically acts as scaffold for the recruitment of DSBs repair proteins at DSB sites. Loss of nuclear TDP-43 as well as ALS-linked TDP-43 mutations are associated with DSB repair defects that trigger neuronal apoptosis in ALS.^40-42^ Moreover, genomic instability due to DNA repair defects and DSBs are also seen in C9orf72-ALS.^52^ Thus, our findings are consistent with these previous reports, indicating that loss of TDP-43 physiological function as a scaffold for DSBs repair proteins due to cytoplasmic TDP-43 redistribution is associated with genomic damage and cytotoxicity in cerebral organoids.

There are some limitations to this study. We used only two cell lines of human iPSCs to create recipient cerebral organoids. Additional iPSCs lines, especially some derived from ALS cases, are required to provide a better understanding of pathogenic TDP-43 propagation in cerebral organoids and possible host factors. Moreover, given that ALS is a late-onset disease, our cerebral organoids at around 10-20 weeks growth *in vitro* do not contain as many mature neurons as an adult human. The ideal cerebral organoids would need to have achieved the same level of maturity as adult CNS tissue, an objective that has yet to be achieved, would be preferable to verify pathogenic TDP-43 propagation. We used cerebral organoids as a CNS tissue model resembling the human brain, while another critical site for TDP-43 pathology in ALS is the spinal cord. In addition, microglia derived from mesoderm, which is an important factor involved in ALS pathophysiology, is deficient in cerebral organoids composed only of cells differentiated from ectoderm. However, robust protocols for creating well-structured spinal cord organoids harboring microglia have not yet been established. It is necessary to wait for such protocols that create organoids that reproduce the anatomical structures and physiological functions of human spinal cord tissue to verify our results.

In conclusion, our findings provide evidence that pathogenic TDP-43 in ALS spinal cord tissue has the potential to spread cell-to-cell to induce TDP-43 pathology as well as astrogliosis and to cause cell cytotoxicity due to DNA damage in human CNS tissue. Although additional investigation is required to elucidate the key factors that induce pathogenic TDP-43 transmission, our model could be used to test treatment strategies aimed at blocking of pathogenic TDP-43 propagation.

## Abbreviations

ALS: amyotrophic lateral sclerosis
DSB: DNA double-strand break
Ipsc: induced pluripotent stem cell
pTDP-43: phosphorylated TDP-43

## Acknowledgements

We are grateful to Dr. Claire Troakes for kindly providing postmortem human tissue samples from the London Neurodegenerative Diseases Brain Bank. We thank the patients and their families for their contributions.

## Supplementary material

Supplementary material is available at *Brain* online.

