## Supplemental material for "Spinal cord extracts of amyotrophic lateral sclerosis spread TDP-43 pathology in cerebral organoids"

| Cell line | Diagnosis | Genetics | Material source | Gender | Age | Ethnicity |
|---|---|---|---|---|---|---|
| AJC001 | Healthy control | N.A. | PBMCs | Male | 37 | Caucasian |
| TD17 | Sporadic ALS-FTLD | Negative for *C9orf72* gene and ALS panel | PBMCs | Female | 65 | Caucasian |

**Supplementary Table 1. Human iPSCs lines information used in this study.** Human iPSCs used in this study were generated from peripheral blood mononuclear cells (PBMCs) of a healthy control (AJC001 line) or sporadic ALS-FTLD patient (TD17 line). N.A. indicates “not assessed”.

| Case number | Diagnosis | *C9orf72* gene | Region | Gender | Age at onset | Age at death | Post-mortem delay (hours) | Source |
|---|---|---|---|---|---|---|---|---|
| Control 1 | Cerebral hemorrhage | N.A. | Cervical | Female | 65 | 65 | unknown | MNI/DBCBB |
| Control 2 | - | - | Cervical | Female | - | 72 | 41 | LNDBB |
| Control 3 | - | - | Cervical | Male | - | 63 | 23 | LNDBB |
| Patient 1 | Sporadic ALS | Normal | Cervical | Male | 46 | 50 | 12 | MNI/DBCBB |
| Patient 2 | Sporadic ALS | Normal | Cervical | Female | 59 | 62 | 22 | MNI/DBCBB |
| Patient 3 | Sporadic ALS | Expanded | Cervical | Male | unknown | 51 | 10 | MNI/DBCBB |
| Patient 4 | Sporadic ALS | Normal | Cervical | Female | unknown | 58 | 31 | MNI/DBCBB |
| Patient 5 | Sporadic ALS | Normal | Cervical | Male | 78 | 79 | 27 | MNI/DBCBB |

**Supplementary Table 2. Human tissue information used in this study.** Postmortem frozen spinal cords tissue specimens were collected from the Montreal Neurological Institute-Hospital (MNI), the Douglas-Bell Canada Brain Bank (DBCBB) and the London Neurodegenerative Diseases Brain Bank (LNDBB). N.A. indicates “not assessed”.

| Antibody (clone) | Source (catalog number) | Host | IF | IHC | WB |
|---|---|---|---|---|---|
| CTIP2 (25B6) | Abcam (ab18465) | rat | 1:500 | N.A. | N.A. |
| PAX6 (Poly19013) | Bio Legend (901301) | rabbit | 1:100 | N.A. | N.A. |
| SOX2 (9-9-3) | Abcam (ab79351) | mouse | 1:200 | N.A. | N.A. |
| TUJ1 (Poly18020) | Bio Legend (802001) | rabbit | 1:5000 | N.A. | N.A. |
| TUJ1 | Abcam (ab41489) | chicken | 1:1000 | N.A. | N.A. |
| phospho TDP-43 (pS409/410) | Cosmo Bio (TIP-PTD-M01) | mouse | 1:1000 | N.A. | 1:2000 |
| phospho TDP-43 (pS409/410) | Cosmo Bio (TIP-PTD-P02) | rabbit | 1:1000 | N.A. | N.A. |
| phospho TDP-43 (pS409) | Cosmo Bio (TIP-PTD-P03) | rabbit | N.A. | 1:1000 | N.A. |
| TDP-43 | Proteintech (10782-2-AP) | rabbit | 1:200 | N.A. | N.A. |
| GFAP | Invitrogen (PA5-16291) | rabbit | 1:200 | N.A. | 1:2000 |
| GR repeat | Proteintech (23978-1-AP) | rabbit | 1:100 | N.A. | N.A. |
| cleaved caspase-3 (E83-77) | Abcam (ab32042) | rabbit | 1:200 | N.A. | 1:500 |
| γH2AX (pS139) | Novus Biologicals (NB100-384) | rabbit | 1:1000 | N.A. | 1:10000 |
| anti-rat Alexa Fluor 488-conjugated | Invitrogen | goat | 1:1000 | N.A. | N.A. |
| anti-rabbit Alexa Fluor 488-conjugated | Invitrogen | goat | 1:1000 | N.A. | N.A. |
| anti-rabbit Alexa Fluor 555-conjugated | Invitrogen | goat | 1:1000 | N.A. | N.A. |
| anti-mouse Alexa Fluor 488-conjugated | Invitrogen | goat | 1:1000 | N.A. | N.A. |
| anti-mouse Alexa Fluor 555-conjugated | Invitrogen | goat | 1:1000 | N.A. | N.A. |
| anti-chicken Alexa Fluor 488-conjugated | Invitrogen | goat | 1:1000 | N.A. | N.A. |
| anti-rabbit peroxidase-conjugated | Invitrogen | donkey | N.A. | N.A. | 1:10000 |
| anti-mouse peroxidase-conjugated | Invitrogen | donkey | N.A. | N.A. | 1:10000 |

**Supplementary Table 3. Antibody information used in this study.** IF, immunofluorescence; IHC, immunohistochemistry; WB, Western blot. N.A. indicates "not assessed".

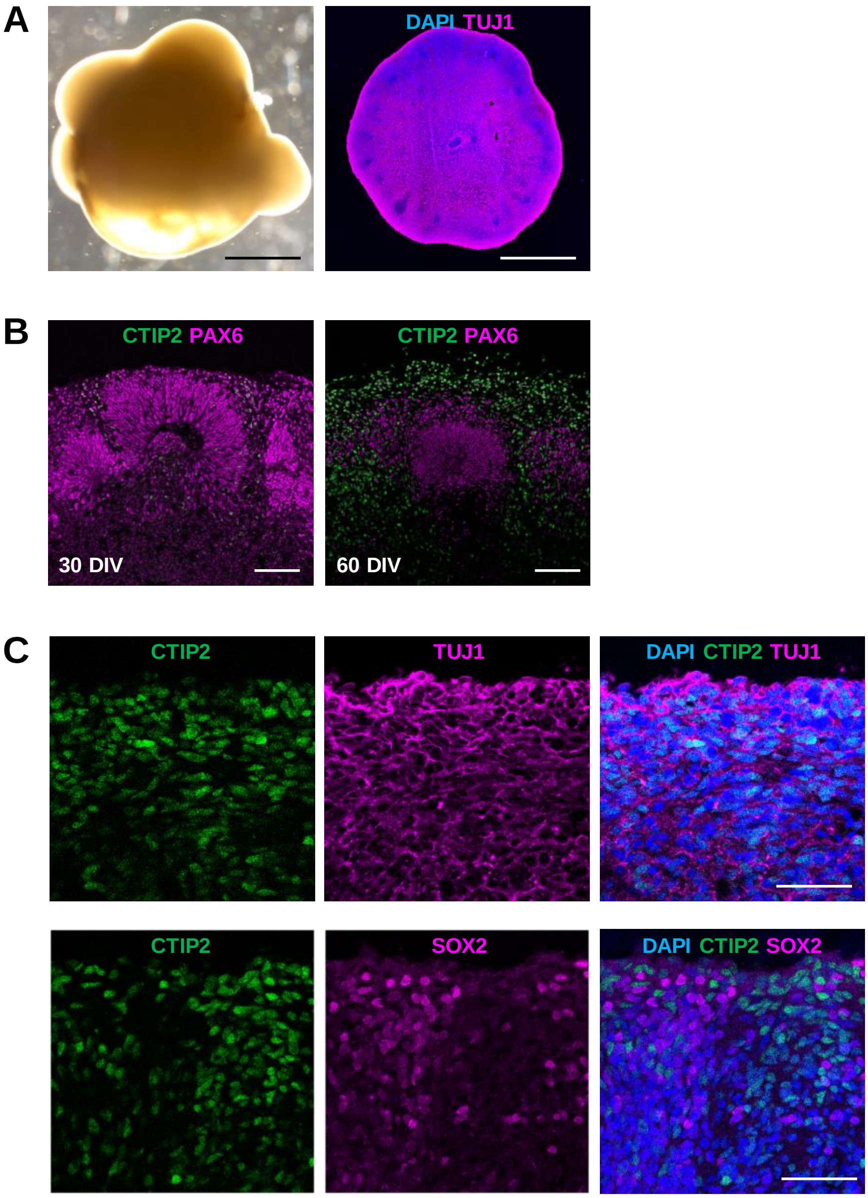


**Supplementary Figure 1. Cerebral organoids exhibit neural cortices-like tissue mimic to human brain.** (**A**) Representative bright-field (*left*) and immunofluorescence images (*right*) of cerebral organoids differentiated from AJC001 iPSCs line at 60 DIV. Scale bars = 1 mm. (**B**) Double-label immunofluorescence images of CTIP2 and PAX6 staining AJC001 cerebral organoids at 30 DIV (*left*) and day 60 DIV (*right*). Scale bars = 100 µm. (**C**) Double-label immunofluorescence images of CTIP2 and TUJ1 (*upper panels*) or SOX2 (*lower panels*) staining AJC001 cerebral organoids at 60 DIV. Sections were counterstained with DAPI to label the nuclei. Scale bars = 50 µm.

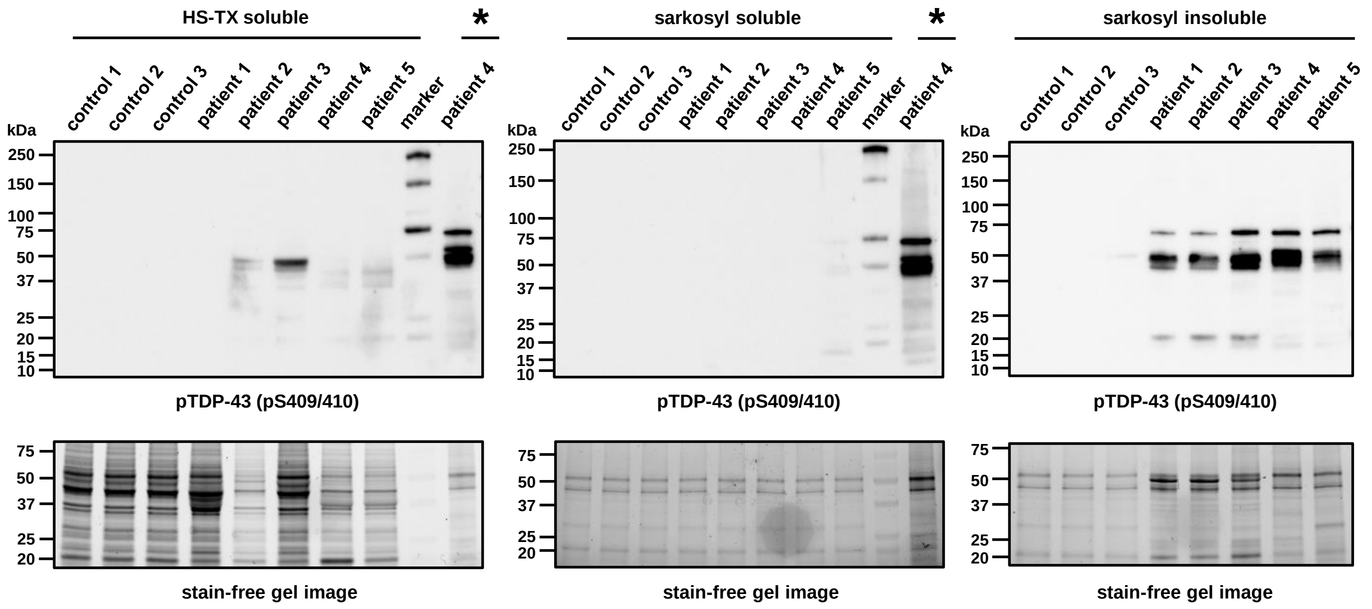


**Supplementary Figure 2. Protein extracts from sporadic ALS spinal cords contain detergent-insoluble pTDP-43.** Western blot analysis of protein extracts from three non-ALS controls (control 1-3) and five sporadic ALS cervical spinal cords (patient 1-5). Postmortem frozen spinal cords were separated into detergent-soluble or -insoluble fractions using high-salt buffer containing 1% Triton-X100 (HS-TX) or 2% sarkosyl buffer. pTDP-43 (pS409/410) antibody was used to detect the detergent-insoluble pathogenic TDP-43. Stain-free gel images were used for protein loading controls. Sarkosyl-insoluble fraction from patient 4 wad loaded on the lanes marked with asterixis (*) as a positive control for pTDP-43 immunoblots.

**A** **B**

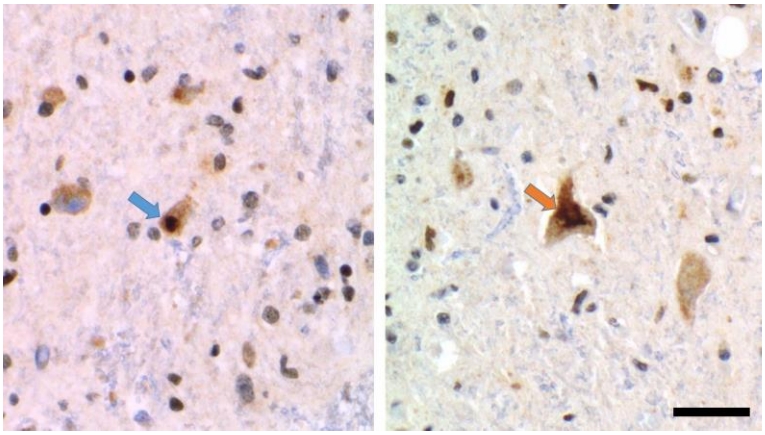

**Supplementary Figure 3. Sporadic ALS-FTLD patient of TD17 iPSCs line displays TDP-43 pathology in anterior horn spinal cords.** (**A and B**) Immunohistochemistry of anterior horn spinal cords stained by TDP-43 antibody. The bule arrow shows punctate intraneuronal inclusion (**A**) and the orange arrow shows skein-like intraneuronal inclusion (**B**). Scale bar = 50 µm.

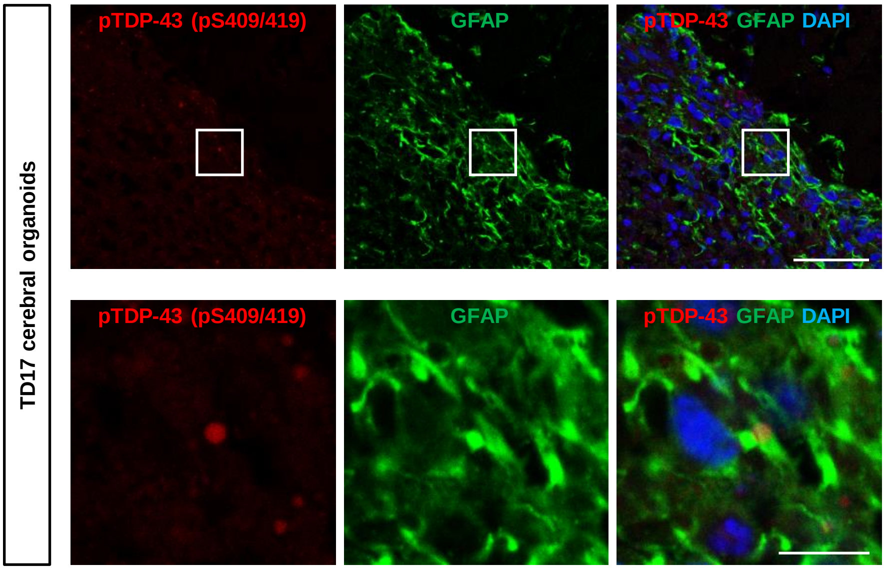


**Supplementary Figure 4. pTDP-43 aggregates distribute in astrocytes.** Immunofluorescence images of TD17 cerebral organoids double-labelled with pTDP-43 and GFAP at 8 weeks post injection of protein extracts from ALS (patient 5). The lower panels are higher magnifications of the white-line boxes in the upper panels. Scale bars = 50 µm (*upper panel*) and 10 µm (*lower panel*).

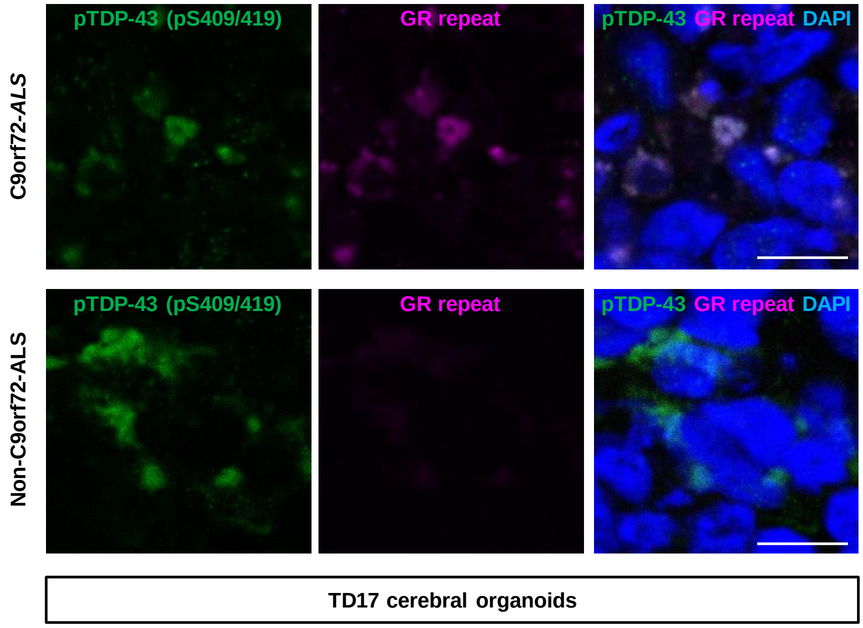


**Supplementary Figure 5. pTDP-43 aggregates colocalize with GR repeat.**
Immunofluorescence images of TD17 cerebral organoids double-labelled with pTDP-43 and GR repeat proteins at 8 weeks post injection of protein extracts from C9orf72-ALS (patient 3) (*upper panels*) or non-C9orf72-ALS (patient 4) (*lower panels*). Scale bars = 10 µm.
